# *In vivo* amyloid-like fibrils produced under stress

**DOI:** 10.1101/2021.02.02.429251

**Authors:** Natália A. Fontana, Ariane D. Rosse, Anthony Watts, Paulo S. R. Coelho, Antonio J. Costa-Filho

**Affiliations:** Departamento de Física, Faculdade de Filosofia, Ciências e Letras de Ribeirão Preto, Universidade de São Paulo, Ribeirão Preto, SP, Brazil; Department of Biochemistry, University of Oxford, Oxford, UK; Departamento de Biologia Celular e Molecular e Bioagentes Patogênicos, Faculdade de Medicina de Ribeirão Preto, Universidade de São Paulo, Ribeirão Preto, SP, Brazil

**Author notes:** Corresponding author: Departamento de Física, Faculdade de Filosofia, Ciências e Letras de Ribeirão Preto, Universidade de São Paulo, Ribeirão Preto, SP, 14040-901, Brazil.

**Keywords:** *in vivo* fibrillation, fluorescence lifetime imaging, Golgi Reassembly and Stacking Proteins, unconventional protein secretion, cellular stress

## Abstract

The participation of amyloids in neurodegenerative diseases and functional processes has triggered the quest for methods allowing their direct detection *in vivo*. Despite the plethora of data, those methods are still lacking. The autofluorescence from the extended β-sheets of amyloids is here used to follow fibrillation of *S. cerevisiae* Golgi Reassembly and Stacking Protein (Grh1). Grh1 has been implicated in starvation-triggered unconventional protein secretion (UPS), and here its participation also in heat shock response (HSR) is suggested. Fluorescence Lifetime Imaging (FLIM) is used to detect fibril autofluorescence in cells (*E. coli* and yeast) under stress (starvation and higher temperature). The formation of Grh1 large complexes under stress is further supported by size exclusion chromatography and ultracentrifugation. The data show for the first time *in vivo* detection of amyloids without the use of extrinsic probes as well as bring new perspectives on the participation of Grh1 in UPS and HSR.

## Introduction

Amyloid fibrils have been a subject of significant interest over the years due to their pivotal participation in several neurodegenerative diseases^[1]^ and, more recently, functional processes.^[2]^ Although the structural 3D arrangement of the fibrils and factors governing their formation have been thoroughly investigated,^[3],4]^ detecting amyloid-like fibrils *in vivo* is still not trivial. Aggregation has been monitored inside cells using GFP-labelled proteins combined with fluorescence methods.^[5]^ The visualization of fibrils has been achieved using strategies based on dyes that bind to amyloid plaques in excised brain tissue,^[6]^ and *in vivo* diagnostics of neurodegenerative diseases have been reported using Magnetic Resonance Imaging.^[7]^ To directly observe fibrils inside cells, other biochemical and biophysical approaches are still necessary.

Recently, Pinotsi *et al.* showed that the formation of amyloid fibrils by tau protein and lysozyme exhibited a characteristic fluorescence in the visible range.^[8]^ The origin of such autofluorescence was attributed to the absorption/emission of electrons delocalized after the formation of hydrogen bonds in the typical β-sheet structure of amyloids, thus allowing low-energy electronic transitions to occur.^[8],9]^ The specific molecular origin of the phenomenon is, however, still not completely understood.^[9]^ Despite this uncertainty, the observed autofluorescence has been firmly correlated with the amyloid formation in several cases, such as Amyloid-β^[10]^ and α-synuclein.^[11]^ Nonetheless, fibril formation monitoring was restricted to the *in vitro* assembly^[10]^ or *in vivo* detection using FRET between an extrinsic probe attached to the protein and the fibril.^[11]^

The use of autofluorescence for direct detection of *in vivo* fibrillation without using an extrinsic probe has not been satisfactorily explored. One reason could be the apparent lack of specificity in detecting the fibril signal due to the competing autofluorescence from the cells. Here, we report results on the *in vivo* formation of amyloid-like fibrils by one member of the Golgi Reassembly and Stacking Protein (GRASP) family without using an extrinsic dye for protein tagging.

GRASPs were initially implicated as participants in the structural organization of the Golgi apparatus,^[12]^ a central organelle in the conventional endoplasmic reticulum (ER)-to-Golgi pathway of protein secretion.^[13]^ However, proteins can reach the plasma membrane and leave the cell via other mechanisms. Unconventional Protein Secretion (UPS) comprises alternatives through which (1) leaderless proteins (lacking the signal sequence for ER localization) are secreted and (2) proteins that use the conventional secretory pathway take a different route traversing from the ER straight to the plasma membrane.^[14],[15]^ Different types of UPS routes have been reported, each dealing with different types of stress.^[14],[15]^ Among the four types of UPS reported thus far, Types III and IV share the common participation of GRASPs.^[16][18]^ In particular, Type III UPS is characterized by the formation of a new GRASP-rich organelle, named Compartment for Unconventional Protein Secretion (CUPS),^[19]^ which leads the secretory cargo to the plasma membrane, where vesicle fuses, releasing its content.

Our group has been exploring the biophysics of GRASPs in the last few years.^[20, 21]^ In one of our previous contributions,^[21]^ we described novel structural features of Grh1, the GRASP from *Saccharomyces cerevisiae*. We have demonstrated that Grh1 contains regions of intrinsic disorder, which seems to be a common feature among GRASPs. Furthermore, it was shown that Grh1 formed amyloid-like fibrils *in vitro*, and the fibrillation was independent of its C-terminal domain.^[21]^ *In vitro* fibrillation has also been observed for both human GRASPs and seems to be another general feature within the GRASP family.^[22],[23]^ A comprehensive review of the biophysics of GRASPs has been recently published.^[24]^

The capacity of Grh1 to form amyloid fibrils is still of unclear biological significance. The presence of functional amyloid aggregates in yeast has been reported previously.^[2]^ We hypothesized that the amyloid-like form of Grh1 also occurs *in vivo*, and the ensemble formed is closely related to the function of Grh1 in UPS, particularly in Type III during starvation,^[25]^ as well as upon increase in temperature (i.e., in Heat Shock Response - HSR).^[26]^ In the case of starvation, intracellular pH drops and becomes acidic,^[27]^ a condition that has been seen to trigger fibrillation *in vitro*. As for the temperature, we have recently reported *in vitro* fibrillation of Grh1 at temperatures greater than 37°C.^[21]^ In the case of yeast, the optimal temperature for its growth is 30°C.^[26]^ The cell can handle mild temperature increases (for instance, from 37 to 41°C) by using the HSR. This is a coordinated event that arrests cell growth through the aggregation of other proteins, thus impairing their function. This aggregation leads to the formation of Stress Granules (SG), which are disassembled by Hsp when thermal stress ceases, and the cell returns to normal growth.^[26]^

Here, we address one issue, namely Grh1 fibrillation *in vivo*, whose contribution is two-fold. On the one hand, the novel direct detection of amyloid fibril formation *in vivo*. On the other hand, the demonstration of the Grh1 fibrillation inside the cell raises new insights towards better understanding fundamental aspects of UPS and HSR in yeast.

## Results

### Grh1 forms in vitro amyloid fibrils under different conditions

The formation of amyloid fibrils *in vitro* has been demonstrated for Grh1.^[21]^ In that study, it was shown that Grh1 fibrillates when submitted either to temperatures higher than 37 °C or changes in the dielectric constant of the medium.^[21]^ Those were two parameters intended to mimic the Grh1 environment (or changes to it) in the cell, i.e., heat shock response and the presence of the membrane field.

Here we complement that study by demonstrating that Grh1 also fibrillates when in acidic pH (< 5.5), a condition that has been shown to happen during starvation. We followed a protocol similar to the one previously used^[8]^ based on the autofluorescence of the fibrils detected using FLIM. The fluorescence decay times were also measured and used to distinguish changes in the fibril formation. Figure S1 shows that Grh1 indeed forms fibrils in the tested conditions. Moreover, the structures formed upon heating or in acidic pH have different fluorescence time decays (the decay is slower in acidic pH), suggesting differences in the fibril structure or the microenvironment where the fibrils are formed. Therefore, the pH-induced *in vitro* fibrillation reinforces our hypothesis of fibrillation in starving conditions in yeast.

### Grh1 is capable of fibrillating in a cellular environment

One of the main challenges in monitoring the fibrillation of a specific protein inside the cell is to find a suitable experimental method to detect fibril formation under the conditions of interest. Fluorescence Lifetime Imaging (FLIM) is based on measuring the time decay of the fluorescence signal after excitation at a determined wavelength and was the method utilized to detect the autofluorescence of fibrils in previous reports.^[8–11]^

In our case, after showing that Grh1 can form amyloid fibrils *in vitro* under conditions that correspond to stress scenarios in the cell, the natural step after that was to ask whether Grh1 could also form fibrils within the cell. Hence, following the same expression protocol described elsewhere,^[21]^ we initially turned to the heterologous expression of Grh1 in *E. coli* to place Grh1 in a cellular environment. Here, the idea was to perform experiments as if we had exchanged the *in vitro* buffer used before for the *E. coli* cytoplasm.

Figure 1 shows the FLIM results obtained from *E. coli* cells at room temperature (Figure 1A), at 37°C (Figures 1B-C), and pH 4.6 (Figure 1D). Using 20 mM sodium acetate in the LB medium as described by Wilks and Slonczewski,^[28]^ we prevented the bacteria from regulating its internal pH, thus turning a decrease in LB pH value into a reduction of the pH of the cytoplasm. The green background in the images is due to the medium autofluorescence. Rod-like structures (colored in green and blue in Figure 1, where colors refer to lifetime values) resolved from the background autofluorescence and likely from Grh1 appeared in the cell cultures containing Grh1 heated at 37°C or in acidic pH (Figures 1C-D). We discarded random fibrillation due to protein overexpression by monitoring a negative control (Figure 1B), where the same *E. coli* strain overexpressing the non-amyloidogenic protein acyl-CoA binding protein (ACBP) did not show significant autofluorescence above the background signal. Other controls in acidic pH did not show any signal either. Therefore, in the conditions that trigger *in vitro* fibrillation (increased temperature or acidic pH), Grh1 also fibrillated inside *E. coli*.

**Figure 1:**
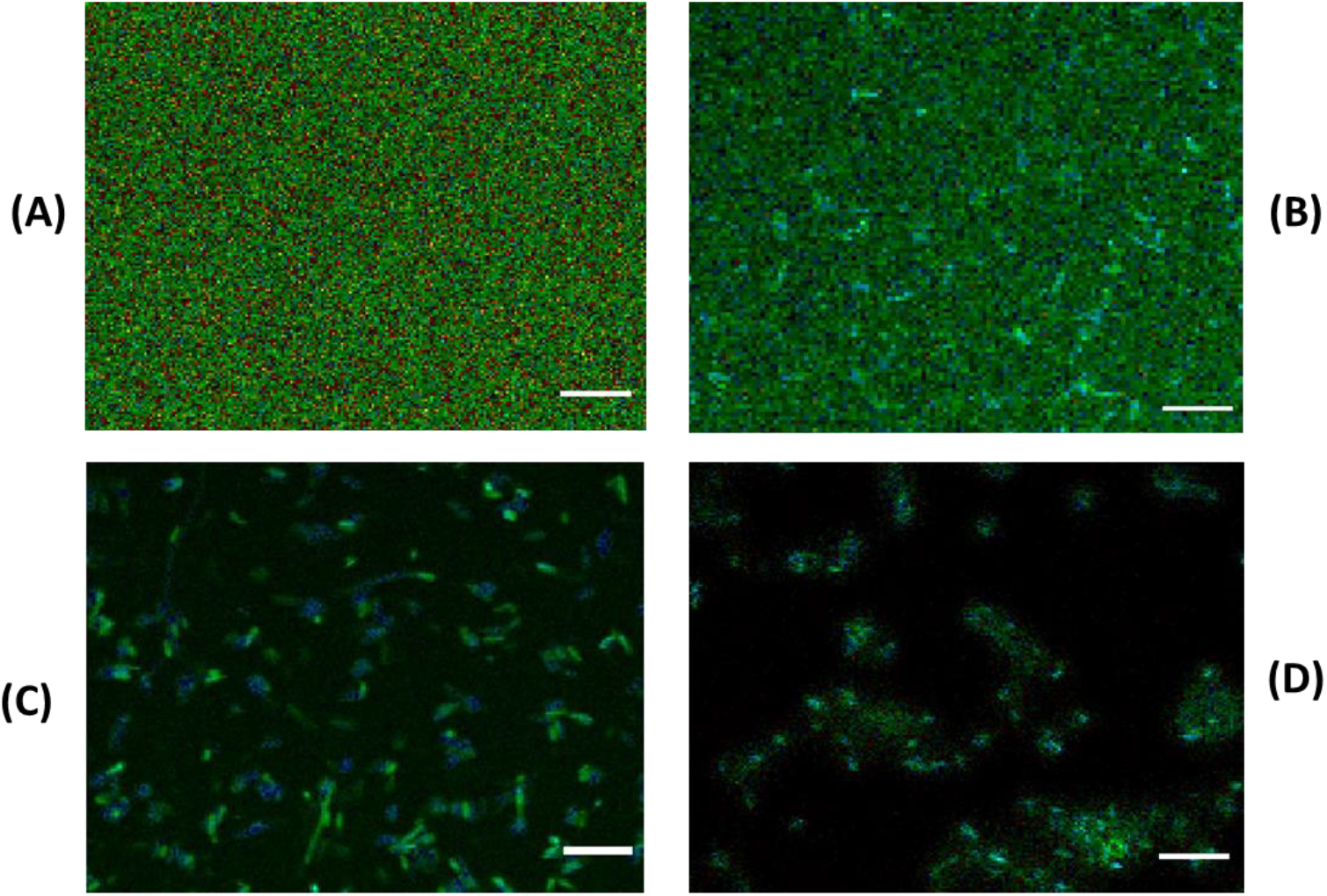
FLIM images of *E. coli* cells excited at 375 nm. The images show *E. coli* cells expressing: (A) Grh1 at room temperature. (B) ACBP in cells heated at 37°C after protein expression. (C) Grh1 in cells heated at 37°C after protein expression. (D) Grh1 in cells in an acidic medium (pH 4.6). Scale bar: 10 μm.

### Fibrillation in yeast under stress

*In cell* Grh1 fibrillation was also investigated in its native environment in *S. cerevisiae* again via FLIM experiments. It is suggested that fibrillation of Grh1 is related to stress, and therefore conditions previously described to trigger Type III UPS^[19]^ and HSR in yeast cells^[27]^ were tested. Figure 2 shows images obtained from a yeast strain called Y270, which does not carry any mutations in *grh1* or related genes and is hereafter referred to as Wild-type (WT) yeast. As expected, there is no autofluorescence in the control cells (Figure 2C). Fluorescence is only observed when the cells are either heated at 37 °C (Figure 2A) or submitted to starvation (Figure 2B), conditions known to trigger HSR or Type III UPS,^[19,27]^ respectively.

**Figure 2:**
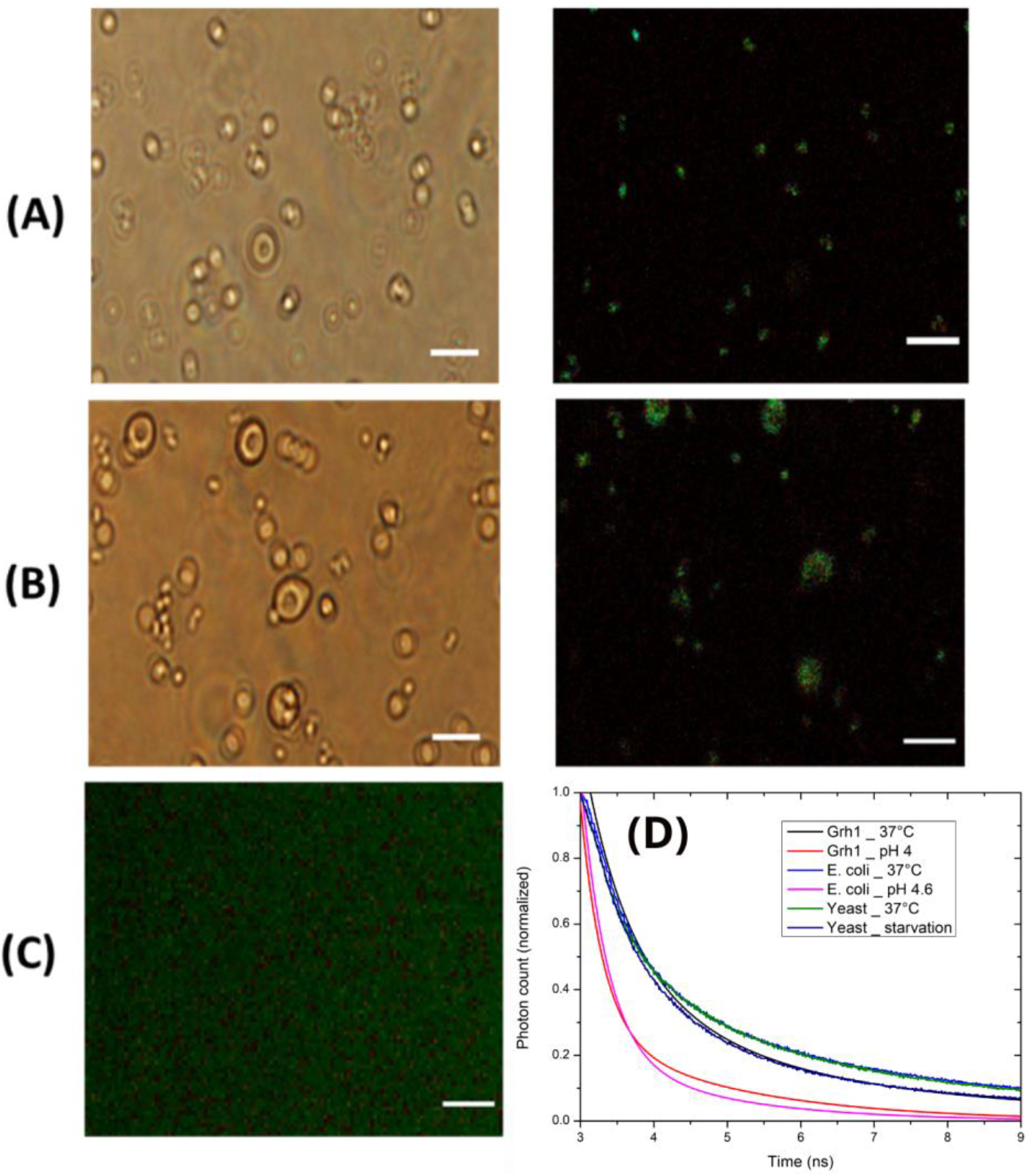
Yeast cell images in the following conditions: (A) heated to 37°C, and (B) under starvation for at least 30 minutes (left: bright field image; right: FLIM image). (C) FLIM image of the control sample. (D) Time decays of fluorescence for purified Grh1 heated at 37°C (black), purified Grh1 in pH 4 (red), *E. coli* at 37°C (light blue), *E. coli* in pH 4.6 (pink), WT yeast at 37°C (green) and starved WT yeast (dark blue). Scale bar: 10 μm.

The time decays of the fluorescence in yeast under starvation or heated at 37°C were measured and compared with those from *E. coli* overexpressing Grh1 and *in vitro* experiments (Figure 2D). The differences between *in cell* and *in vitro* time decays of fluorescence are expected since lifetime depends on the fluorophore environment.^[29]^ In Figure 2D, we can see that each type of stress yielded different time decays, which suggests differences in the environment of the fibrils upon temperature increase, starvation, or acidic pH. This is an interesting finding since variations in the cell responses to those stress conditions are expected.^[19],[26]^ Moreover, the temperature stress in both cell types (*E. coli* and yeast) led to identical time decays, which were somewhat longer than the time decays observed in the other two stress conditions.

### The lack of Grh1 changes the autofluorescence signal

The results shown in Figures 1 and 2 indicate *in cell* fibrillation of Grh1. Still, it could be argued that the fluorescence exhibited was due to other fibrillation processes taking place inside the yeast. Similar experiments were carried out to better understand the origin of the observed fibrillation process, but this time using a Grh1 knockout yeast lineage.

As expected, under physiological conditions, no autofluorescence was observed (data not shown). However, in the sample heated at 37°C (Figure 3A), there was a change in the fluorescence pattern. Not only was the FLIM signal from Grh1 lost, but also the cells appeared as black dots (Figure 3A – highlighted by the red circles in the right panel). Although it was not possible to quantitatively determine the loss of signal from this experiment only, the events occurring in the cell suggest considerable changes took place during HSR when Grh1 was not present.

**Figure 3:**
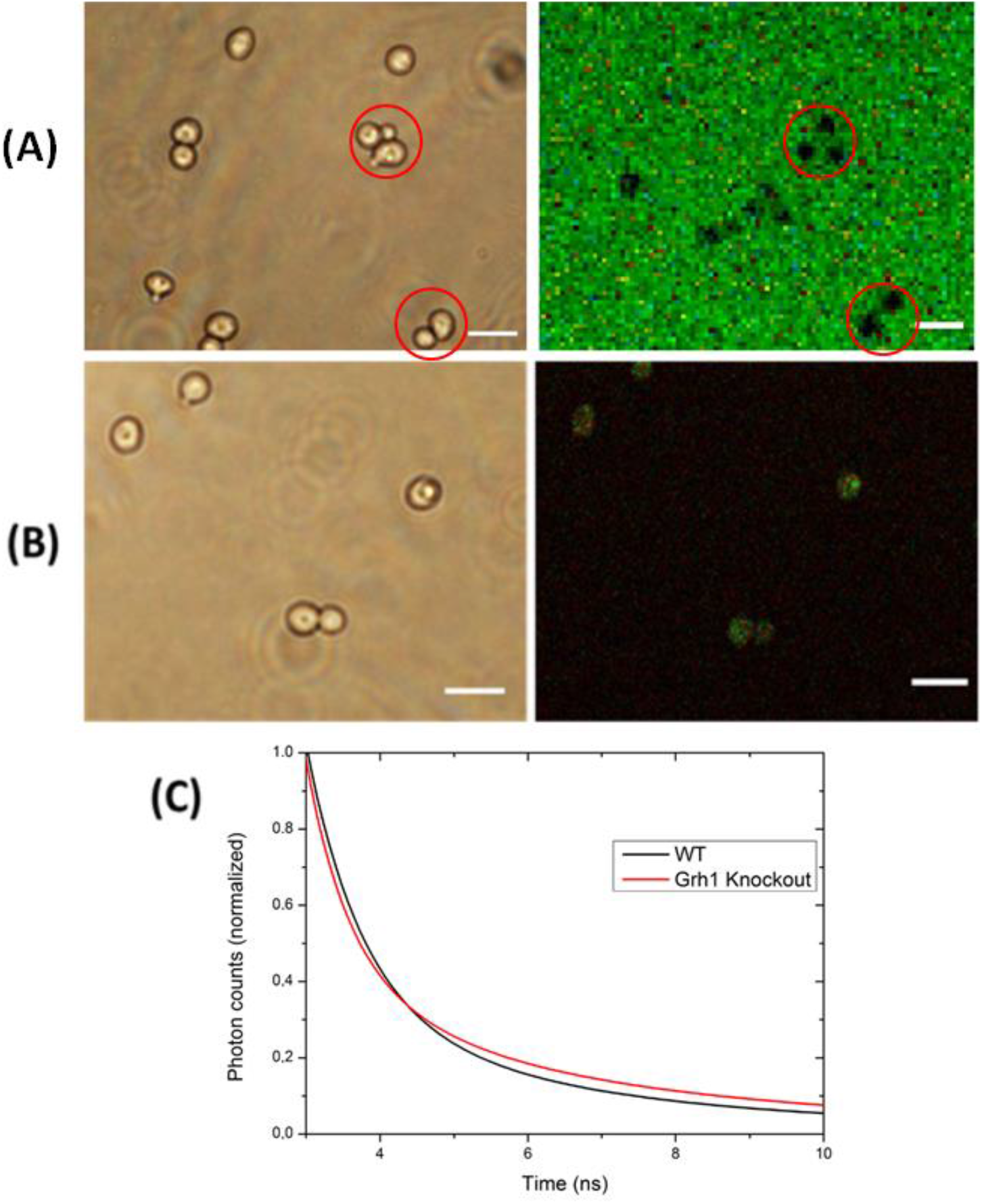
Images of Grh1 knockout yeast cells in the following conditions: (A) heated at 37° C and (B) under starvation for at least 30 minutes (left: bright field image; right: FLIM image). (C) Time decays of fluorescence for WT (black) and Grh1 knockout (red) cells. Scale bar: 10 μm.

In starvation, some autofluorescence in the Grh1-knockout cells (Figure 3B) was observed. To investigate this somewhat unexpected result, we also examined the pattern seen in the lifetime values by looking at the distribution of those time decays obtained from WT and Grh1-knockout cells (Figure S2). A shift to longer lifetime values was observed in the distribution from the Grh1-knockout cells compared with the WT sample. The lifetime corresponding to the peak of the distribution for Grh1-knockout cells increased, and the highest value of the observed lifetimes was longer (around 7.5 ns for WT cells compared to ca. 10 ns for knockout cells). The decay times of fluorescence in both samples (Figure 3C) were also different [(4.60±0.02) for WT cells, and (5.60±0.05) for knockout cells], which confirms the contribution (or the absence of it) of Grh1 to the detected signal. An explanation for those results would be that the presence of Grh1 in starvation conditions would give rise to a dominant fibrillation process, thus sequestering the FLIM signal and leading to a more Gaussian-like shape of the respective lifetime distribution.

To further investigate the fibrillation in starvation conditions, we used a GFP-tagged yeast strain and the immunoprecipitation of Grh1-GFP from cultures either grown in optimal conditions (control) or starved for 2 hours. Once control and test group cells were grown, they were lysed, and an immunoprecipitation protocol followed as described in the Material and Methods. The FLIM experiments were then performed as before, but instead of whole cells, the samples, in this case, consisted of trapped GFP (and, consequently, Grh1). The results for the control and starvation conditions can be seen in Figure S3. While there was no signal coming from the control sample (Figure S3A), in the immunoprecipitated sample from starved cells (Figure S3B), the signal presented a decay of (4.8±0.1) ns that is very similar to the (4.60±0.02) ns lifetime obtained from the whole cells in starved conditions (Figure S3C). The slightly slower decay might be explained by the reorganization of the fibrils once released from the cell’s interior.

### Grh1 forms higher-order complexes in certain conditions

A different strategy to test the formation of large complexes is size exclusion chromatography (SEC), in which large complexes (in our case, fibrils) are excluded from the separation column. To track Grh1 as it moves through the column, we employed the GFP-tagged yeast strain. Yeast cells were cultivated under the desired condition: control, starvation, or submitted to a temperature increase. The whole extract obtained from disrupted yeast cells in each condition was then applied in a SuperDex200, and the proteins were tracked via their optical absorbance at 280 and 395 nm for the target proteins and GFP, respectively. Figure 4A shows the results of the SEC experiments. For clarity, elution profiles were normalized, and only the signals at 395 nm are shown. In the control experiment (black line), the signal from GFP appeared around 16 mL of elution. In starved cells (red line), the curve was shifted to the left, indicating that GFP (and consequently Grh1) was then too large to enter the column, being excluded at 9 mL. On the other hand, there were two populations in the non-permissive temperature condition (37 °C, blue line): one that was excluded from the column, and another one that left the column at the same point as in the control condition. While SEC does not give information about the type of structure formed, it does indicate that, under stress conditions, Grh1 undergoes changes that significantly affect its size.

**Figure 4:**
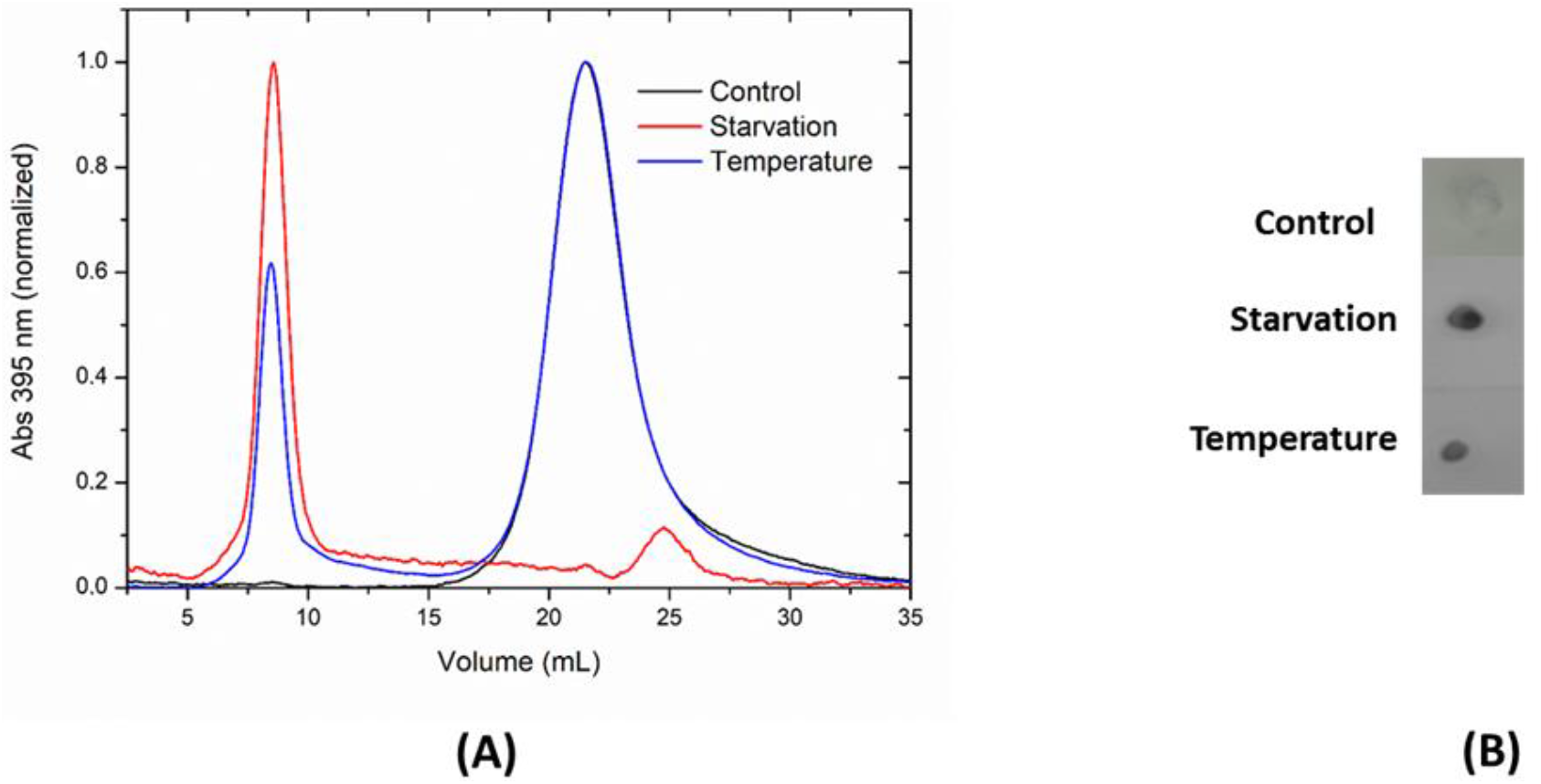
Results of the experiments monitoring the formation of higher-order complexes by using the yeast strain producing a GFP-tagged Grh1. (A) Elution profiles (normalized) of the size exclusion chromatography following the GFP signal (at 395 nm). (B) Dot blot of the pellet obtained from the ultracentrifuged samples. The detection was based on the use of an anti-GFP antibody.

To explore whether the observed size change in Grh1 was due to protein aggregation, ultracentrifugation in SDS was used to differentiate amyloid fibrils from other possible amorphous aggregates of Grh1.^[5]^ We also used the Grh1-GFP strain and performed a dot blot (Figure 4B) to detect the presence of Grh1 in the non-solubilized pellet. Confirming the SEC observations, Grh1-GFP fibrils were present under stress conditions but not in the control, indicating that Grh1 formed SDS-insensitive large complexes when the yeast cells were submitted to starvation or non-permissive temperature. Therefore, we can infer that the size increase was not due to amorphous aggregation but rather to amyloid formation.

### Fibrillation reversibility

After incubation in starvation conditions, yeasts can return to their normal state when physiological conditions are restored.^[25]^ In agreement with that, our data show that there was no autofluorescence from the sample that was subjected to starvation for 2 hours and then brought back to normal conditions (Figure S4A). This suggests that all the fibrillation events indicated by the autofluorescence in starvation conditions were reversible. Moreover, it is known that HSR can sustain the stress for 2 hours and the cells can also go back to normal once the optimal temperature is restored.^[30]^ Akin to starved cells, the autofluorescence from cells subjected to heating at 37°C for 2 hours (heat shock) was no longer seen when the cells were brought back to 30°C (Figure S4B).

### Visualizing Grh1 fibrils

Transmission Electron Microscopy (TEM) was used to visualize the assemblies formed by Grh1 in different conditions: purified Grh1 heated to 37°C for 30 minutes before the preparation of the TEM grids; Grh1-GFP immunoprecipitated from control (optimal growth conditions) cells and cells subjected to heat shock or starvation; pellets of ultracentrifuged WT control cells and cells subjected to starvation. The results of each experiment are shown in Figure 5.

**Figure 5:**
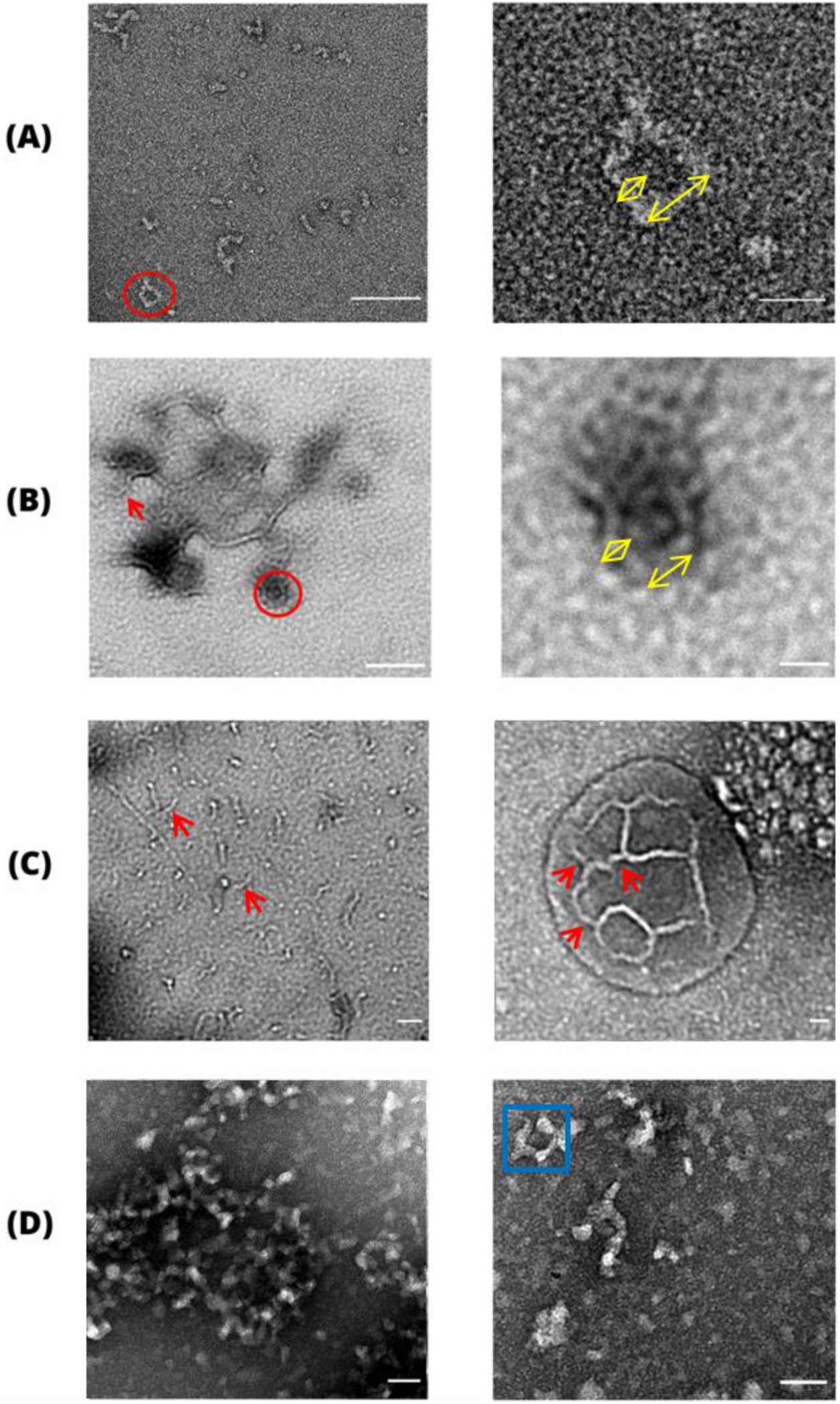
TEM images of: (A) purified Grh1 heated at 37°C. (B) Grh1-GFP immunoprecipitated from yeast cells in heat shock. Right panels are zoomed-in images of the red circle seen on the left panels. Yellow double arrows indicate measured dimensions of particles. (C) Grh1-GFP immunoprecipitated from yeast cells in starvation. Red arrows indicate units of fibrils. (D) Pellets of ultracentrifuged yeast cells in starvation. Blue square highlights the structures found on the grids. Scale bar: (A), (B), (C) left panels:100 nm. (A), (B), (C) right panels, (D): 20 nm.

Samples of purified Grh1 resulted in the images of fibrils shown in the left panel of Figure 5A. Their amyloid signature has been previously confirmed.^[21]^ Curiously, besides the usual fibrillar arrangement, it is also possible to observe the formation of a seemingly closed square-like structure (red circle in the left panel of Figure 5A). The dimensions of the particles in the grids (both the ones forming the square-like structure and the loose ones) were obtained by using the software ImageJ (see yellow double arrows in the right panel of Figure 5A), yielding lengths between 15 and 20 nm, and thickness in the range of 4.0 to 7.5 nm.

In Figure 5B, we have images of immunoprecipitated Grh1-GFP from cells in heat shock. In this case, the structures present in the grid looked more fibrillar than the ones from the *in vitro* experiment (purified Grh1, Figure 5A). Such fibrils seem to be formed by individual units that are linked together to build the complete structure. One of the individual units is marked with a red arrow in the left panel of Figure 5B. The size and thickness of those units are compatible with what was found for purified Grh1 (16 to 21 nm in length and 4.0 to 7.5 nm in thickness). It is possible again to find arrangements of the fibrils consisting of square-like structures (zoomed-in in the right panel of Figure 5B). For the sample immunoprecipitated from cells subjected to starvation, we obtained the images in Figure 5C. Akin to the heat-shock condition, we have fibril-like assemblies. It is also possible to identify small individual units (marked with red arrows in the left panel of Figure 5C). These have sizes and thickness similar to those found in the heat-shock condition’s grids (Figure 5B). Unlike the previous conditions, starved cells also gave rise to a structurally different arrangement (shown in the right panel of Figure 5C) obtained from another grid of the same experiment. In this case, even though it was difficult to differentiate all the individual units that make up the whole structure, some of them could be distinguished (see red arrows), forming a more complex structural arrangement than those seen in Figures 5A-B. Such arrangement resembled a network of branched convoluted tubules. The dimensions of the individual units were in the same range as before (16-22 nm of length and 4.5-5.8 nm of thickness), and the average diameter of the larger structure was ca. 153 nm.

Considering we know from previous experiments (Figure 4B) that Grh1 was found in the pellet of starved cells subjected to ultracentrifugation, we further investigated the fibrillation in starvation by taking the pellet of starved WT cells to the microscope and obtained images like the ones in Figure 5D. After the ultracentrifugation, the type of arrangement found was slightly different from those seen in the sample obtained using the immunoprecipitation protocol (Figure 5C). That can be due to the physical process of ultracentrifugation or to changes caused by GFP in the structure. Nevertheless, one can again see small units forming interconnected fibril- or square-like structures (left panel in Figure 5D). These squares can be seen in more detail in the right panel of Figure 5D (marked with a blue square).

Interestingly, the dimensions of the squared structures (*i.e.,* the size of their sides) were around 21 nm of side length and 5 nm of thickness. These values are compatible with the ones found in the experiments with purified Grh1 (*in vitro*) and immunoprecipitated samples in both conditions (heat shock and starvation, *in vivo*). The ability to form somewhat short fibrils that can apparently result in square-like structures seems to be the organization pattern adopted by Grh1.

Finally, it is worth mentioning that the grids containing Grh1-GFP immunoprecipitated from cells grown in optimal growth conditions or pellet of ultracentrifuged cells grown in optimal growth conditions neither showed any of the structures described above nor any other of relevance.

## Discussion

Amyloid formation inside cells has been a subject of significant interest over the years. It has gained even more attention lately since various aggregation-prone proteins, whose potential formation of amyloid-like structures is not disease-related, have been described.^[2]^ Nevertheless, detecting amyloid-like fibrils *in vivo* remains a challenge. Several reports have successfully probed aggregation within cells^[5]^ by using chimeras of the target protein tagged with fluorescent reporters. In the case of aggregation-prone proteins, the foci formed inside the cells can, in principle, be visualized under the fluorescence microscope.^[5]^ Despite the usefulness of this latter approach, in the specific case of Grh1, compartmentalization of the protein either in CUPS (in the case of starvation) or likely in stress granules during HSR can lead to the formation of 1 to 3 punctate structures, therefore hampering the direct identification of potential fibrils.

Grh1 fibrillation has been demonstrated *in vitro*^[21]^ and we further expanded the list of triggering factors of that process to now include acidic environments (Figure S1). However, its occurrence within the cell and its potential implications were still unclear. Fibrils and changes in the cytoplasmic state are seemingly tools used by the cell to cope with different types of stress.^[2, 30]^ Our initial hypothesis was then that fibrils of Grh1 could form *in vivo* and would be necessary under specific conditions. The test of our idea was mainly based on the use of a label-free assay, firstly described by Pinotsi *et al.* in 2013.^[8]^ Such a method for detecting amyloid fibrils relies on the intrinsic fluorescence in the UV-visible region that arises when the protein changes conformation and adopts the characteristic β-sheet rich structure of amyloids. Chan *et al.* ^[9]^ further described this fluorescence signature and how unlikely it would be for an experiment based solely on fibril autofluorescence to be used *in vivo* due to the competing autofluorescence from the cells. It was not a surprise, then, those publications using FLIM and intrinsic fluorescence that followed were based on either FRET experiments or changes in a reporter’s physicochemical parameter when fibrillation happened.^[10, 11]^

Therefore, although *in vivo* FLIM of proteins without a reporter may not always be possible, we could show this can be a valuable approach for yeast cells. We specifically tested Grh1 *in vivo* fibrillation due to starvation (Type III trigger – Figures 2B and 3B) or to temperature increase (HSR trigger – Figure 2A), two stress scenarios where Grh1 seems to play a central role. During starvation, Grh1 leaves the Golgi membrane and relocalizes to be part of CUPS.^[16]^ Upon heat shock, although there is hitherto no study describing the direct participation of Grh1, its increased expression when the yeast is subjected to thermal stress has been reported.^[31]^ In control conditions, the autofluorescence signal arising from the cells was negligible when excited at 375 nm (Figure 2C).

Our data showed that the autofluorescence could be used to monitor the formation of the fibrils themselves (presence or absence – Figures 1, 2, and 3) and differences in the environment surrounding them induced by the distinct stress sources (Figure 2D). More specifically, the alterations in pH, temperature, and starvation yielded distinguishable lifetime decays, therefore strongly suggesting that the fibrils are in different environments, which agrees with the idea of distinct cell responses and the corresponding Grh1 function in each case. It is interesting to note that the temperature stress in both cell types (*E. coli* and yeast) led to identical time decays, which indicates that the cell’s HSR involves the formation of the fibrils in somewhat similar environments.

To further infer the participation of Grh1 fibrillation in each stress condition, we obtained FLIM and time decay data from a Grh1-knockout yeast strain under heat shock and starvation (Figure 3). The images from the heat-shocked knockout cells, unlike the images of the heat-shocked Grh1-containing cells (Figure 2C), surprisingly showed the cells as black dots, which indicates the lack of Grh1 led the cells to a different type of response to HS. As previously said, currently, no studies link Grh1 to the HSR. Still, some of the data available and our results reinforce the hypotheses of fibrillation of Grh1 in the context of HS. Gasch *et al.*,^[31]^ through microarray DNA experiments, measured the changes in transcript levels over time in response to several types of stress, including temperature. For Grh1, what they observed was an increase in transcription when the cell was subjected to heat shock (37°C).^[31]^ Considering a stress situation where the cell stops its non-essential activities to save as much energy as possible, an increase in Grh1 transcription suggests a protein function in that scenario. Besides that, when the Grh1-GFP cell is subjected to a non-permissive temperature (37°C), the signal from GFP, which was initially more dispersed, coalesced into foci inside the cell, thus becoming brighter spots than before (Figure S5). As described by Alberti, Halfmann, and Lindquist,^[5]^ proteins that fibrillate *in vivo* coalesce into microscopic assemblies, just like the ones we observed for Grh1.

On the other hand, the knockout cells showed a distinct response to starvation compared to HS. In this case, FLIM signals could be detected (Figure 3B), thus suggesting fibrillation of other proteins in the cell, described before for the protein Cdc19.^[34]^ Despite the existence of this non-Grh1-related signal, we could find differences in the time decays (Figure 3C) and their distributions (Figure S2) measured from WT and knockout cells. Furthermore, sedimentation experiments and a modified version of the filter retardation assay (in the form of an SEC experiment)^[5]^ were used to corroborate our FLIM data by showing that Grh1 was present in the fibrils formed within the yeast cells. To do so, we used a yeast strain expressing Grh1 tagged with GFP to allow for the immunoprecipitation of the Grh1-GFP chimera. The fibrils of Grh1-GFP were then detected using FLIM (Figure S3B) and compared with the data obtained for the whole yeast cell (Figure 2B). The pattern observed in those images and the similar lifetimes measured in both experiments (Figure S3C) allowed us to infer that the *in-cell* fibrillation was indeed due to Grh1.

To enhance our understanding of the fibril morphology, we used transmission electron microscopy to visualize the structures formed by Grh1. The TEM images from purified Grh1, immunoprecipitated Grh1-GFP from cells in heat shock and starvation, and ultracentrifuged WT cells in starvation (Figure 5) all showed similar patterns where one can see the coexistence of fibril-like and square-like structures. The experiments here reported did not allow for atomic-resolution structural determination of Grh1-containing assemblies. Nevertheless, it was possible to distinguish short linear units in some images, whose dimensions are ca. 20 nm long and 6 nm thick. They are apparently linked to form the fibril itself and other more complex arrangements. Bruns *et al.* ^[32]^ showed that, upon starvation, Grh1 concentrates in large membraneous punctae in the cell to form the so-called compartment for unconventional protein secretion (CUPS). Despite the thorough description of the biogenesis of this new compartment, its detailed structure and composition were still not clear. Curwin *et al.* ^[33]^, in an elegant combination of correlative light and electron microscopy (CLEM) and fluorescence, advanced the knowledge on that issue by reporting CUPS were arranged in a somewhat spherical structure of convoluted tubules and vesicles, whose average diameter was ca. 200 nm. In the same paper, the authors proposed a pathway for CUPS formation that would consist of the segregation of Grh1 in tubular clusters, followed by the engulfment of this immature CUPS by a sheet-like structure called saccule. The exact origin of the saccule membrane was not determined, but Grh1-containing membranes were suggested as one possibility. Our TEM data (Figure 5) offer another possibility: the Grh1 higher-order structural arrangements. The GRASP’s ability to form fibrillar structures, Grh1 among them, has been demonstrated.^[21]^ In the Grh1 case, combining individual somewhat linear units seems to give rise to distinct 3D structures, such as the fibrils and the square-like seen in Figures 5B-C. Furthermore, in the right panel of Figure 5C, an even more complex arrangement involving an apparent network of tubules, whose rough average diameter was around 150 nm, can be seen. The dimensions of the chains formed by the units reported here are compatible with the images showing the different stages of CUPS maturation described by Curwin *et al.* ^[33]^ as the saccule. We could speculate that the 3D-sheet of unknown origin engulfing the immature CUPS and even the Grh1-containing vesicles described by Curwin *et al.* could be formed by Grh1 higher-order structures rather than their monomers. The reversibility of Grh1 fibril formation (Figure S4) is also compatible with a mechanism that needs to be turned on and off depending on the conditions triggering the cell stress.

Our data do not rule out the participation of secretory and endosomal membranes, but Grh1 structural plasticity and its ability to form complex arrangements seem to be a new piece of information that needs to be added as an alternative in the formation of compartments for UPS as well as for heat shock response. Furthermore, the capacity of Golgi-related proteins to undergo liquid-liquid phase separation (LLPS) has been recently demonstrated^[34]^, which raises another exciting possibility regarding whether GRASP could also undergo such transition, a feature yet to be determined. LLPS as the source of membrane-less compartments has gained particular interest over the last few years^[35]^, and it might well be another aspect to consider when tackling the problem of UPS and HSR.

In summary, here, we present solid evidence of *in vivo* formation of Grh1 fibrils under stress conditions. While the details on how exactly they function in both Type III UPS and HSR are yet to be fully revealed, our data reinforce the new concept of functional fibrils in yeasts as active factors in response to certain types of stress. As any first-time idea appearing in the literature, our findings seem to bring more questions than answers. Still, they undoubtedly offer new perspectives to better understand amyloids in vivo and Grh1 roles in UPS and HSR.

## Materials and Methods

### Protein Expression and Purification

The protocol for Grh1 expression in *E. coli* and purification is described elsewhere.^[21]^ For the experiments with *E. coli* in acidic pH, cells were grown, and expression was induced and carried out for 18 hours at 20 °C. Cells were pelleted by centrifugation at 8,000g for 10 minutes, transferred for LB medium added of 50 mM MES and 20 mM sodium acetate, pH 4.6, and kept shaking at 20 °C for 1 hour.

### Yeast Culture

The *Saccharomyces cerevisiae* strain Y470 was used in the FLIM experiments with excitation at 375 nm. All the other experiments without tagging or knockout were performed in the parental strain BY4741. The GFP-tagged and the knockout strains come from the commercial Yeast GFP clone collection (ThermoFisher Scientific). All cultures were grown in YPD medium (1% Yeast Extract, 2% Peptone, 2% Glucose).

For all experiments, a colony of the desired strain was placed in liquid YPD or SC-ura and allowed to grow overnight at 30° C. The following morning the colony was transferred to 200 ml of YPD and allowed to grow again, this time under slow agitation until the culture reached an optical density at 600 nm of ca. 0.5.

For starvation, cultures were centrifuged and resuspended in 2% potassium acetate solution (as described by Cruz-Garcia *et al.* ^[25]^). For HSR, cultures were placed under agitation at 37 °C. Both experiments were run for 2 ½ hours, and the same time was used for recovery, where the cultures were put back at optimal growth conditions.

### Immunoprecipitation

Immunoprecipitation was carried out using the GFP-tagged strain with the GFP-Trap Magnetic Agarose (Chromotek). Cells in the desired conditions were centrifuged at 12,000g for 10 minutes, and the supernatant was collected. The beads were previously equilibrated with dilution/washing buffer (10 mM Tris pH 7.5, 150 mM NaCl, 0.5 mM EDTA. pH adjusted to 8). Beads were added to the supernatant, and the solution was left rotating end-to-end for 1 hour at 4°C. Two washing steps were performed by separating the beads with a magnet until the supernatant was clear, discarding the supernatant, and resuspending beads with washing buffer.

For transmission electron microscopy experiments, the proteins were eluted by adding 50μL 0.2 M glycine pH 5.5. Although the protocol recommends pH 2.5, given that Grh1 fibrillates in pH below 5.5, we decided to use this value (5.5) to avoid undesired fibrillation in the samples. The samples were incubated for 30 seconds under constant agitation, the supernatant was placed in another Eppendorf tube, and the solution neutralized with 5 μL 1M Tris base, pH 10.4. For confocal and FLIM experiments, since the presence of the beads was not a problem, the elution step was not performed, and the beads with trapped GFP were taken to the microscope.

### Fluorescence Lifetime Imaging Microscopy

FLIM experiments were performed in an IX71 Inverted Microscope (Olympus) equipped with a PicoQuant MT 200 confocal module. Excitation was set at 375 nm, and emission was detected from 405 nm. Data collection and analysis were performed with the SymPhoTime 64 software (PicoQuant). A Region of Interest (ROI) was delimited for analysis, and the nExponential Tailfit mode was selected. The number of exponential components was chosen based on the best distribution of the residuals and the best x^2^ value found. To obtain the lifetime’s histograms, the FLIM fit function was used based on the manual provided by PicoQuant (available at: https://www.picoquant.com/images/uploads/page/files/17319/4_td_flim.pdf). Decay curves presented are the best fits for each of the results.

### Multiphoton Microscopy

Images were acquired using a LSM 780 Multiphoton AxioObserver (Zeiss), equipped with a titanium sapphire laser. Excitation was set at 880 nm, and emission recorded with a 515/30 nm filter. The microscope was available at the Laboratório Multiusuário de Microscopia Multifóton (Departamento de Biologia Celular e Molecular e Bioagentes Patogênicos, Faculdade de Medicina de Ribeirão Preto, Universidade de São Paulo).

### Gel Filtration

Cultures in different conditions were centrifuged and pelleted in PBS, followed by sonication for 7 minutes, in cycles of 30 sec on and 30 sec off. To separate solid particulates, cultures were centrifuged at 20,000g for 20 minutes, and the collected supernatant was concentrated using a Vivaspin column (GE Healthcare, Buckinghamshire, United Kingdom). The concentrated supernatant was applied to a Superdex200 10/300 GL gel filtration column (GE Healthcare, Buckinghamshire, United Kingdom).

### Ultracentrifugation

2 mL of cultured cells were harvested and resuspended in 300 uL of lysis buffer (50 mM Tris, pH 7.5, 150 mM NaCl, 2 mM EDTA, 1mM PMSF, 5% glycerol). Disruption was carried out by vortexing with acid-washed glass beads. 300 μL of RIPA buffer (50 mM Tris, pH 7.0, 150 mM NaCl, 1% Triton X-100, 0.1% SDS) were added to the solution, which was vortexed again for 10 sec. Centrifugation at 1,500g for 5 minutes was used to pellet cell debris. 200 μL of the soluble fraction was centrifuged in a TLA 100-2 rotor for 45 min at 100,000g and 4 °C in an Optima TL Beckman ultracentrifuge.

### Dot Blot

10 μL of resuspended precipitate (see Ultracentrifugation) was blotted in nitrocellulose membrane, which was then blocked with Blocking buffer (TBST - 10 mM Tris, 150 mM NaCl, 0.1% Tween 20-plus 1% w/v BSA) for 24 hours. Incubation with rabbit-polyclonal anti-GFP primary antibody in a 1:10,000 dilution (Invitrogen Cat. Number #A-11122) in Blocking Buffer was carried out for one hour, followed by three washes with TBST. Incubation with anti-Rabbit Secondary Antibody Solution Alk-Phos. Conjugated (Invitrogen™) followed and preceded for one hour. Washing and detection were performed as recommended for Novex® AP Chromogenic Substrate.

### Transmission Electron Microscopy

The purified protein was analyzed at the Brazilian Nanotechnology National Laboratory in a JEOL 3010. The ultracentrifuged and immunoprecipitated samples were analyzed in an FEI Tecnai 12 Transmission Electron Microscope at the Sir William Dunn School of Pathology of the University of Oxford (UK). A 120 kV of acceleration voltage was applied on the samples deposited in a 15 mA discharged copper grid and stained with 2% uranyl acetate. The images were analyzed with ImageJ.^[36]^

## Author Contributions

NAF performed most of the experimental procedures and analyses. ADR was involved in part of the experimental procedures. AW and PSRC coordinated part of the project. AJCF conceived and coordinated the project. All authors also approved the final version of the manuscript.

## Acknowledgments

The authors acknowledge Fundação de Amparo à Pesquisa do Estado de São Paulo (FAPESP) for the financial support (Grant no. 2015/50366-7, 2012/20367-3 and 2008/55831-6). NAF and ADR thank FAPESP for the Ph.D. and undergraduate fellowships (Grant no. 2016/23863-2 and 2018/24968-8). AJCF thanks Conselho Nacional de Desenvolvimento Científico e Tecnlógico (CNPq) for the partial financial support (Grant no. 306682/2018-4). The authors also thank Prof. Cleslei Fernando Zanelli (Department of Biological Sciences, School of Pharmacy, UNESP, Brazil) for the Grh1 GFP-tagged and Grh1 knockout yeast strains.

## References

[1] E. H. Koo, P. T. Lansbury, J. W. Kelly, Proceedings of the National Academy of Sciences of the United States of America 1999, 96 (18), 9989, https://doi.org/10.1073/pnas.96.18.9989.

[2] G. Cereghetti, S. Saad, R. Dechant, M. Peter, Cell Cycle 2018, 17 (13), 1545, https://doi.org/10.1080/15384101.2018.1480220.

[3] M. G. Iadanza, M. P. Jackson, E. W. Hewitt, N. A. Ranson, S. E. Radford, Nature Reviews Molecular Cell Biology 2018, 19 (12), 755, https://doi.org/10.1038/s41580-018-0060-8.

[4] Z. L. Almeida, R. M. M. Brito, Molecules 2020, 25 (5), https://doi.org/10.3390/molecules25051195.

[5] S. Alberti, R. Halfmann, S. Lindquist, Methods in Enzymology, Vol 470: Guide to Yeast Genetics:: Functional Genomics, Proteomics, and Other Systems Analysis, 2nd Edition 2010, 470, 709, https://doi.org/10.1016/s0076-6879(10)70030-6.

[6] M. T. Elghetany, A. Saleem, Stain Technology 1988, 63 (4), 201, https://doi.org/10.3109/10520298809107185.

[7] E. M. Reiman, W. J. Jagust, Neuroimage 2012, 61 (2), 505, https://doi.org/10.1016/j.neuroimage.2011.11.075.

[8] D. Pinotsi, A. K. Buell, C. M. Dobson, G. S. K. Schierle, C. F. Kaminski, Chembiochem 2013, 14 (7), 846, https://doi.org/10.1002/cbic.201300103.

[9] F. T. S. Chan, G. S. K. Schierle, J. R. Kumita, C. W. Bertoncini, C. M. Dobson, C. F. Kaminski, Analyst 2013, 138 (7), 2156, https://doi.org/10.1039/c3an36798c.

[10] E. K. Esbjorner, F. Chan, E. Rees, M. Erdelyi, L. M. Luheshi, C. W. Bertoncini, C. F. Kaminski, C. M. Dobson, G. S. K. Schierle, Chemistry & Biology 2014, 21 (6), 732, https://doi.org/10.1016/j.chembiol.2014.03.014.

[11] G. S. K. Schierle, C. W. Bertoncini, F. T. S. Chan, A. T. van der Goot, S. Schwedler, J. Skepper, S. Schlachter, T. van Ham, A. Esposito, J. R. Kumita, E. A. A. Nollen, C. M. Dobson, C. F. Kaminski, Chemphyschem 2011, 12 (3), 673, https://doi.org/10.1002/cphc.201000996.

[12] F. A. Barr, M. Puype, J. Vandekerckhove, G. Warren, Cell 1997, 91 (2), 253, https://doi.org/10.1016/s0092-8674(00)80407-9.

[13] C. Viotti, in Unconventional Protein Secretion, Humana Press, New York 2016.

[14] C. Rabouille, V. Malhotra, W. Nickel, Journal of Cell Science 2012, 125 (22), 5251, https://doi.org/10.1242/jcs.103630.

[15] W. Nickel, C. Rabouille, Nature Reviews Molecular Cell Biology 2009, 10 (2), 148, https://doi.org/10.1038/nrm2617.

[16] F. Giuliani, A. Grieve, C. Rabouille, Current Opinion in Cell Biology 2011, 23 (4), 498, https://doi.org/10.1016/j.ceb.2011.04.005.

[17] D. Brough, P. Pelegrin, W. Nickel, Journal of Cell Science 2017, 130 (19), 3197, https://doi.org/10.1242/jcs.204206.

[18] V. Malhotra, Embo Journal 2013, 32 (12), 1660, https://doi.org/10.1038/emboj.2013.104.

[19] D. Cruz-Garcia, V. Malhotra, A. J. Curwin, Seminars in Cell & Developmental Biology 2018, 83, 22, https://doi.org/10.1016/j.semcdb.2018.02.021.

[20] L. F. S. Mendes, A. F. Garcia, P. S. Kumagai, F. R. de Morais, F. A. Melo, L. Kmetzsch, M. H. Vainstein, M. L. Rodrigues, A. J. Costa, Scientific Reports 2016, 6, https://doi.org/10.1038/srep29976; L. F. S. Mendes, L. G. M. Basso, P. S. Kumagai, R. Fonseca-Maldonado, A. J. Costa, Biochimica Et Biophysica Acta-General Subjects 2018, 1862 (4), 855, https://doi.org/10.1016/j.bbagen.2018.01.009; L. F. S. Mendes, N. A. Fontana, C. G. Oliveira, M. Freire, J. L. S. Lopes, F. A. Melo, A. J. Costa-Filho, Febs Journal 2019, 286 (17), 3340, https://doi.org/10.1111/febs.14869; S. T. Reddy, L. F. S. Mendes, N. A. Fontana, A. J. Costa, International Journal of Biological Macromolecules 2019, 135, 481, https://doi.org/10.1016/j.ijbiomac.2019.05.089; L. F. S. Mendes, M. R. B. Batista, P. J. Judge, A. Watts, C. Redfield, A. J. Costa, Febs Journal 2020, 287 (15), 3255, https://doi.org/10.1111/febs.15206.

[21] N. A. Fontana, R. Fonseca-Maldonado, L. F. S. Mendes, L. P. Meleiro, A. J. Costa, Scientific Reports 2018, 8, https://doi.org/10.1038/s41598-018-33955-1.

[22] S. T. Reddy, V. N. Uversky, A. J. Costa-Filho, Int J Biol Macromol 2020, 162, 1982, https://doi.org/10.1016/j.ijbiomac.2020.08.126.

[23] T. S. Reddy, V. N. Uversky, A. J. Costa-Filho, Eur Biophys J. 2020, 49(2), 133, https://doi.org/10.1007/s00249-019-01419-7.

[24] L. F. S. Mendes, N. A. Fontana, S. T. Reddy, V. N. Uversky, A. J. Costa-Filho, Int J Biol Macromol 2020, https://doi.org/10.1016/j.ijbiomac.2020.08.203.

[25] D. Cruz-Garcia, A. J. Curwin, J. F. Popoff, C. Bruns, J. M. Duran, V. Malhotra, Journal of Cell Biology 2014, 207 (6), 695, https://doi.org/10.1083/jcb.201407119.

[26] S. Lindquist, Annual Review of Biochemistry 1986, 55, 1151, https://doi.org/10.1146/annurev.bi.55.070186.005443.

[27] R. Orij, J. Postmus, A. Ter Beek, S. Brul, G. J. Smits, Microbiology (Reading) 2009, 155 (Pt 1), 268, https://doi.org/10.1099/mic.0.022038-0.

[28] J. C. Wilks, J. L. Slonczewski, J Bacteriol 2007, 189 (15), 5601, https://doi.org/10.1128/JB.00615-07.

[29] A. Jain, C. Blum, V. Subramaniam, in (Ed.: Elsevier), 2009, Ch. 4.

[30] S. Saad, G. Cereghetti, Y. Feng, P. Picotti, M. Peter, R. Dechant, Nat Cell Biol 2017, 19 (10), 1202, https://doi.org/10.1038/ncb3600; E. W. Wallace, J. L. Kear-Scott, E. V. Pilipenko, M. H. Schwartz, P. R. Laskowski, A. E. Rojek, C. D. Katanski, J. A. Riback, M. F. Dion, A. M. Franks, E. M. Airoldi, T. Pan, B. A. Budnik, D. A. Drummond, Cell 2015, 162 (6), 1286, https://doi.org/10.1016/j.cell.2015.08.041.

[31] A. P. Gasch, P. T. Spellman, C. M. Kao, O. Carmel-Harel, M. B. Eisen, G. Storz, D. Botstein, P. O. Brown, Mol Biol Cell 2000, 11 (12), 4241, https://doi.org/10.1091/mbc.11.12.4241.

[32] C. Bruns, J. M. McCaffery, A. J. Curwin, J. M. Duran, V. Malhotra, J Cell Biol 2011, 195 (6), 979, https://doi.org/10.1083/jcb.201106098.

[33] A. J. Curwin, N. Brouwers, M. Alonso Y Adell, D. Teis, G. Turacchio, S. Parashuraman, P. Ronchi, V. Malhotra, Elife 2016, 5, https://doi.org/10.7554/eLife.16299.

[34] P. Ziltener, A. A. Rebane, M. Graham, A. M. Ernst, J. E. Rothman, FEBS Lett 2020, 594 (19), 3086, https://doi.org/10.1002/1873-3468.13884.

[35] S. Boeynaems, S. Alberti, N. L. Fawzi, T. Mittag, M. Polymenidou, F. Rousseau, J. Schymkowitz, J. Shorter, B. Wolozin, L. Van Den Bosch, P. Tompa, M. Fuxreiter, Trends Cell Biol 2018, 28 (6), 420, https://doi.org/10.1016/j.tcb.2018.02.004.

[36] C. T. Rueden, J. Schindelin, M. C. Hiner, B. E. DeZonia, A. E. Walter, E. T. Arena, K. W. Eliceiri, BMC Bioinformatics 2017, 18 (1), 529, https://doi.org/10.1186/s12859-017-1934-z.

