## Supplementary figures for "*In vivo* amyloid-like fibrils produced under stress"

**Supplementary Material**

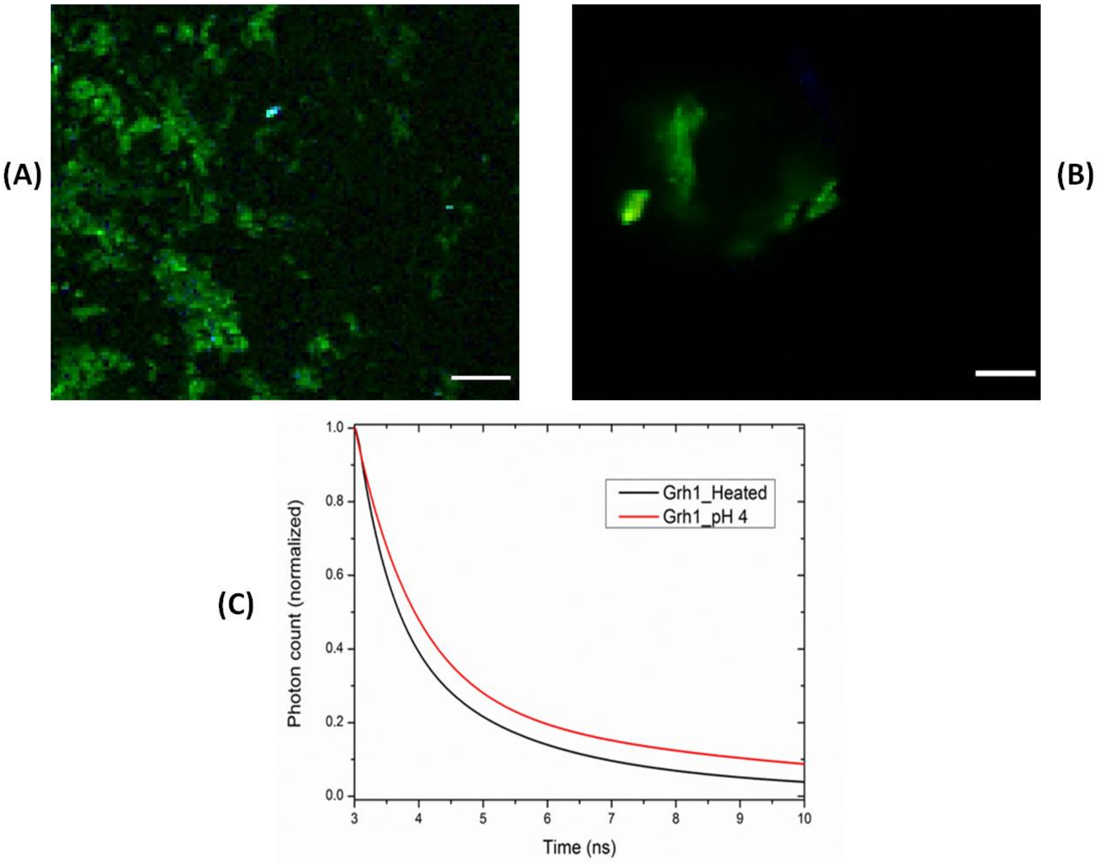


**Figure S1:** *In vitro* fibrillation of Grh1 in different conditions. FLIM images of purified Grh1 excited at 375 nm: (A) heated to 37°C, (B) at pH 4, (C) time decays of fluorescence for heated Grh1 (black) and at pH 4 (red). Scale bar: 10 µm.

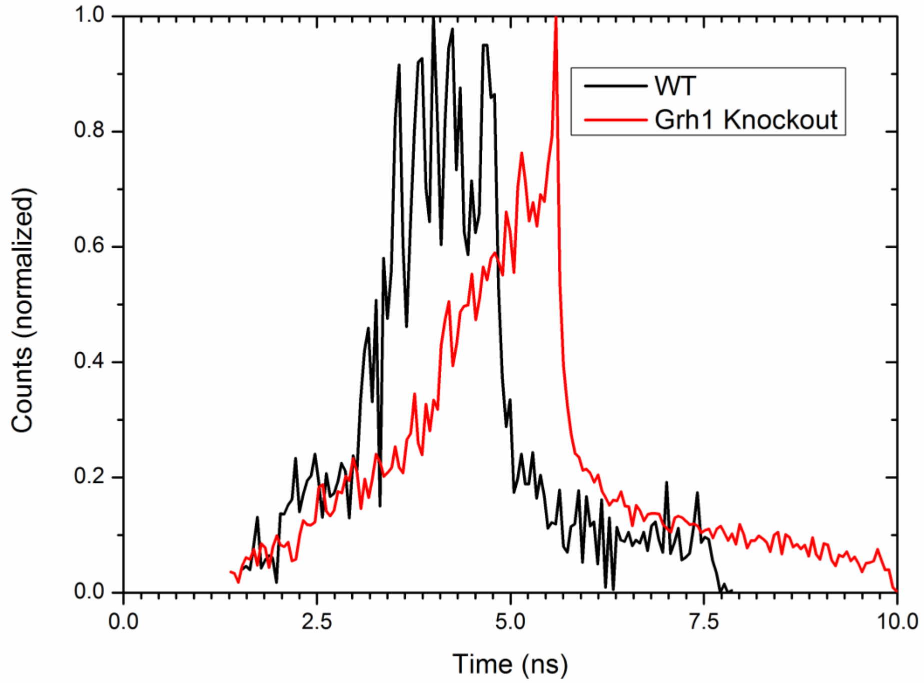


**Figure S2:** Histograms of lifetime values obtained from WT (black) and Grh1-knockout (red) yeast cells subjected to starvation and excited at 375 nm.

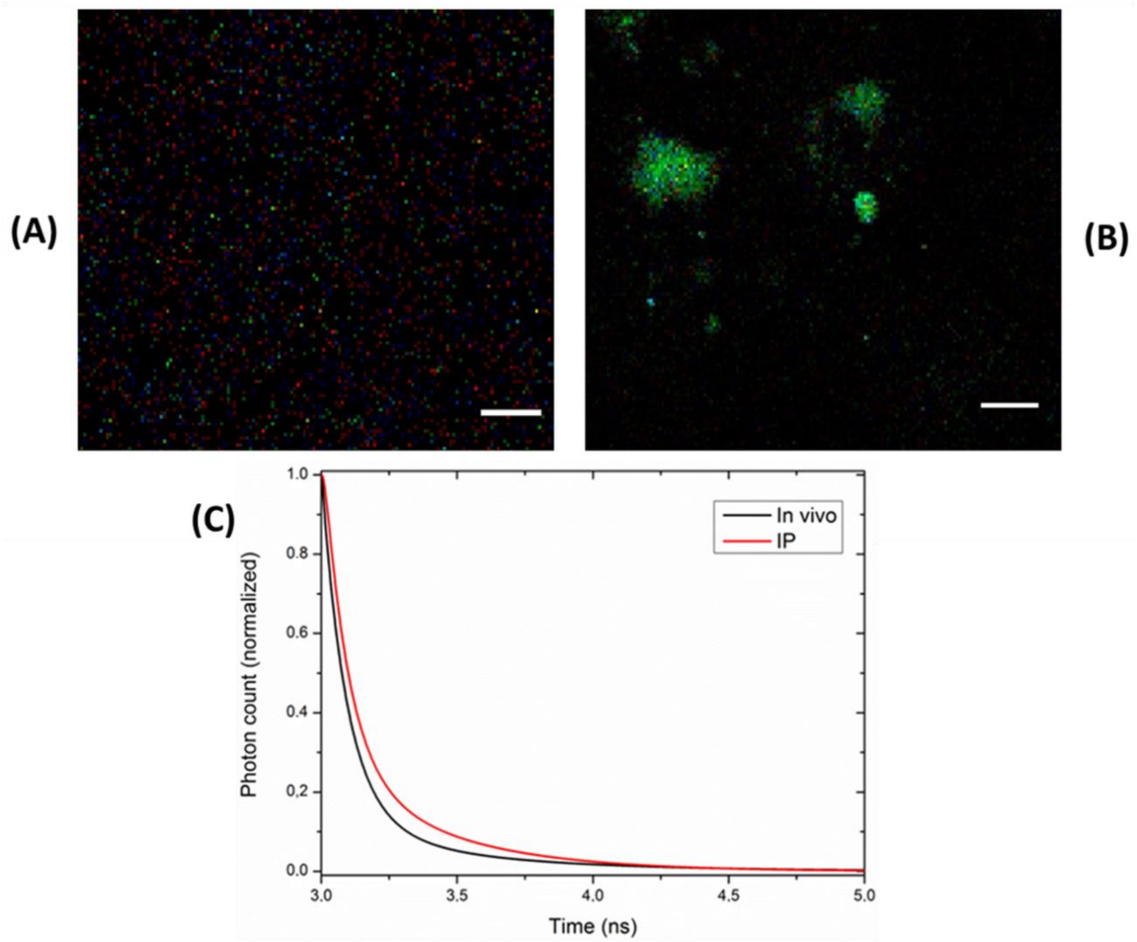


**Figure S3:** Results of the experiments with GFP-tagged Grh1: (A) FLIM image of the control sample, (B) FLIM image of the fibrils immunoprecipitated from cells submitted to starvation, and (C) time decays of fluorescence from the cells in starvation (black) and from the immunoprecipitated sample (IP - red). Scale bar: 10 µm.

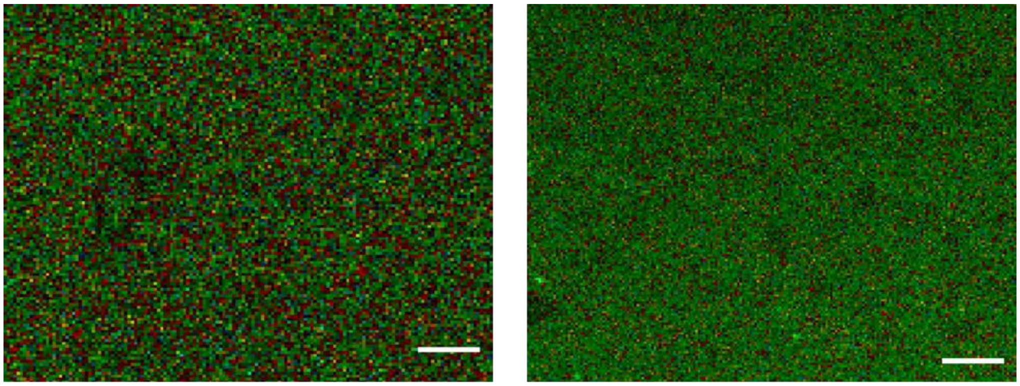

(A) (B)

**Figure S4:** Reversibility experiments. FLIM images of cells submitted to (A) starvation, and (B) heat-shock and cycled back to non-stress conditions. Scale bar: 10 µm.

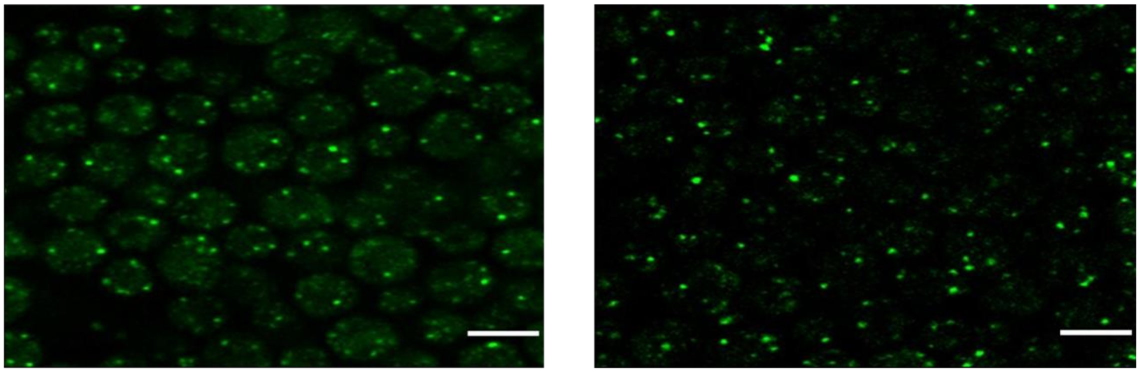

(A) (B)

**Figure S5:** Images from multiphoton microscopy of GFP-tagged Grh1 cells excited at 880 nm: (A) control; (B) heated at 37°C. Scale bar: 5 µm
